# Re-engineering of TNF_α_-NF-_κ_B signalling dynamics in cancer cells using pathogenic *E. coli* effectors

**DOI:** 10.1101/2023.03.03.530985

**Authors:** Qiyun Zhong, Francesca Butera, Gad Frankel, Chris Bakal

**Affiliations:** Dynamical Cell Systems, Chester Beatty Laboratories, The Institute of Cancer Research, London, United Kingdom; Department of Life Sciences, Imperial College, London, United Kingdom

## Abstract

Re-engineering NF-κB signalling towards enhancing beneficial outcomes such as tumour cell elimination, while minimising inflammatory damage, is a potential therapeutic avenue. In this study, we explored the ability of bacterial effectors injected into host cells by the type III secretion system to regulate NF-κB translocation dynamics. We used the enteropathogenic *Escherichia coli* effectors Tir (NF-κB activator), NleC (NF-κB protease) and NleE (TAB2/3 methyltransferase), to manipulate NF-κB translocation and cancer cell survival. We discovered that while these effectors have either limited or no cytotoxicity alone, they greatly enhanced caspase-8-dependent pancreatic cancer cell death in the presence of TNFα. Single cell analysis revealed that the sub-population of cells showing high NF-κB activation is less susceptible to cell death caused by NleC or NleE but instead is more susceptible to Tir. A combination of Tir, NleE and TNFα eliminated 95% cancer cells with limited NF-κB activation, potentially due to NleE-dependent blockage of the immediate pro-survival NF-κB activation without inhibiting Tir’s long-term NF-κB activation that promotes cell death. This work demonstrates that effector combinations could be used to re-engineer stress responses towards favourable outcomes.

## 1. Introduction

The transcription factor NF-κB is the main regulator of immune responses, inflammation and cell fate. In the absence of stimulation, NF-κB is sequestered in the cytosol by inhibitor of κB (IκB). External stimuli, such as the cytokine tumour necrosis factor α (TNFα) or the bacterial lipopolysaccharide (LPS), trigger signalling pathways that converge on the TAB/TAK1 complex-mediated phosphorylation of IκB kinase (IKK), followed by IκB phosphorylation and removal, freeing the canonical NF-κB dimer RelA/p50 which subsequently translocates into the nucleus to initiate gene expression. NF-κB target genes promote protective immunity against pathogens and/or anti-tumorigenic activity; for example, engaging in immune cell activation, antigen presentation and cytokine production. NF-κB transcribes genes encoding cytokines, chemokines, immunoreceptors, growth factors as well as both activators and inhibitors for various cell death types (Bauernfeind et al., 2009; Hinz et al., 1999; Kreuz et al., 2001; Liu et al., 2017; Micheau et al., 2001; Schwenzer et al., 1999; Stehlik et al., 1998; Wang et al., 1998, 1999; Yang et al., 2015). The oscillation of NF-κB between nuclear and cytosol, is dependent on stimuli type, dosage and timing as well as cell type (Butera et al., 2022; Nelson et al., 2004); leading to different transcriptional outcomes (Colombo et al., 2018; Tian et al., 2005; Zambrano et al., 2016; Zhao et al., 2018).

Under classical stimuli (e.g. TNFα), NF-κB promotes the rapid transcription of anti-apoptotic genes including TRAF2, cIAPs, cFLIP and Bcl-2 family proteins (Kreuz et al., 2001; Schwenzer et al., 1999; Stehlik et al., 1998; Tian et al., 2005; Wang et al., 1998, 1999; Zhao et al., 2018) but also apoptosis genes FAS and FASL and, under atypical stimuli (e.g. UV-C and anthracyclines), represses XIAP and Bcl-XL to promote apoptosis (Campbell et al., 2004; Ho et al., 2005; Liu et al., 2012; Singh et al., 2007; Zheng et al., 2001). NF-κB activation is also involved in p53- and reactive oxygen species-dependent apoptosis (Ricca et al., 2001; Ryan et al., 2000). Moreover, NF-κB stimulates the expression of NLRP3 and caspase-4, priming the cells for inflammasome formation and the pro-inflammatory cell death pyroptosis (Bauernfeind et al., 2009; Yang et al., 2015). Therefore, depending on the context of activation, NF-κB can promote inflammation and effects as diverse as cell death versus survival.

The diverse effects of NF-κB signalling are exemplified by the dual roles of TNFα and NF-κB in tumorigenesis. TNFα was initially discovered to induce haemorrhagic necrosis in tumour. It is one of the main anti-tumour strategies of immune cells (Kearney et al., 2018). However, its endogenous level is believed to be usually insufficient in tumour suppression without additional sensitisation using chemotherapy drugs (Habtetsion et al., 2018). Mild NF-κB activation is required for the tumour-suppressing activity of multiple chemotherapy drugs. However, extensive NF-κB activation induced by chemotherapy drugs is associated with tumour metastasis and drug resistance (Behranvand et al., 2022; Vyas et al., 2014). Furthermore, the accumulation of TNFα produced by tumour-associated macrophages and the resulting chronic activation of NF-κB promote tumour proliferation, vascularisation and/or chemoresistance, which are frequently detected in the tumour microenvironment and associated with poor prognosis of many cancer types, notably pancreatic ductal adenocarcinoma (PDAC) which has the highest mortality of all cancers (Laha et al., 2021; Montfort et al., 2019; Weichert et al., 2007; Zhao et al., 2016). While TNFα treatment enhances the growth and metastasis of PDAC via NF-κB activation, TNFα blockade (e.g. etanercept) reduces the activation of NF-κB and the production of pro-inflammatory proteins such as IL-8, resulting in an inhibition of the growth and metastasis of implanted PDAC in mice, but failed to increase survival in PDAC clinical trials (Egberts et al., 2008; Herman et al., 2013; Padoan et al., 2019; Wu et al., 2013; Zhao et al., 2016). Long-term use of anti-TNFα therapy can also lead to an increased risk of infections and even cancer, potentially due to the importance of TNFα in the immune system (Li et al., 2021). In addition, although numerous small molecule drugs have been found to have NF-κB regulating abilities with potential therapeutic implications, they face challenges of target specificity, adverse drug reactions and drug resistance (Perkins and Gilmore, 2006; Ramadass et al., 2020). It is therefore important to expand our search for alternative methods to manipulate the TNFα-NF-κB axis. One therapeutic avenue may be to develop methods that enhance the anti-tumorigenic activity of TNFα, while limiting the pro-inflammatory and pro-survival effects of NF-κB.

Apart from the traditional small molecule drugs and antibody-derived drugs, experimental therapeutics have explored the possibility of microbial toxins in cancer treatment (Kramer et al., 2018; Rüter and Schmidt, 2017; Zahaf and Schmidt, 2017). A group of small bacterial proteins translocated into the host cytosol by the bacterial surface syringe-like nanomachine type III secretion system (T3SS), called T3SS effectors, have interesting properties with potential therapeutic values such as the ability to cooperate with each other or with host cytokines to generate different downstream outcomes and broad-spectrum targeting of common pathways in immune responses (Rüter and Schmidt, 2017; Shenoy et al., 2018). NF-κB, owing to its extensive involvement in host immune responses, is a major target of T3SS effectors.

Human diarrhoeal & tumour-colonising pathogen enteropathogenic *Escherichia coli* (EPEC) uses an array of T3SS effectors as an interactive network to manipulate NF-κB signalling and the downstream cell survival (Levine et al., 1978; Magdy et al., 2015; Shenoy et al., 2018). The most rapidly translocated effector, translocated intimin receptor (Tir), is mainly responsible for binding to the bacterial surface protein intimin, which promotes intimate bacterial attachment and actin polymerisation at the infection site (Gruenheid et al., 2001; Kenny et al., 1997; Nougayrède et al., 2003). Independent of its actin modulation activity, Tir also promotes the activation of NF-κB and caspase-4/GSDMD-dependent pyroptosis in colorectal cancer cells (Zhong et al., 2020, 2022). In turn, seven effectors target different positions of the NF-κB signalling pathway to dampen immune responses. As NF-κB signalling pathway is interwoven with cell death pathways, effectors acting upstream on the TNF receptor complexes, NleB1/2 and EspL, can inhibit both TNFα-induced NF-κB activation as well as extrinsic apoptosis or necroptosis (Giogha et al., 2021; Li et al., 2013; Pearson et al., 2017) and the multifunctional effectors, NleH1/2, can target intrinsic apoptosis independent of their regulatory functions on the NF-κB co-factor RPS3 (Gao et al., 2009; Royan et al., 2010). Others, NleC (broad-spectrum NF-κB protease) and NleE (methyltransferase targeting TAB2/3 to prevent IKK function), are relatively specific to NF-κB inhibition as they have no known direct involvement in cell death (Baruch et al., 2010; Nadler et al., 2010; Newton et al., 2010; Yen et al., 2010; Zhang et al., 2012). Although the target proteins of individual effectors have largely been discovered, how effectors cooperate to regulate single cell NF-κB signalling dynamics leading to different cell fate outcomes is poorly characterised.

We aimed to determine if NF-κB signalling in cancer cells could be re-engineered by EPEC effector combinations to dictate NF-κB signalling dynamics and NF-κB-regulated cell fates – i.e. cell survival vs cell death. Such work sets the basis for identifying combinations which may have therapeutic value by promoting cell death while limiting damaging inflammatory responses. Toward this goal we used high-content single cell live imaging combined with cell death analysis to explore the potential use of individual effectors and effector combinations to regulate the TNFα- NF-κB signalling pathway to generate different outcomes. We show that the NF-κB signalling pathway activator Tir as well as the NF-κB signalling pathway inhibitors NleC and NleE can all promote TNFα-dependent cell death, despite their differential regulation of NF-κB signalling pathway; while Tir and NleC has antagonistic effects on cell death, combining Tir, NleE and TNFα could further promote cell death to eliminate almost the entire cancer cell population but with limited NF-κB activation.

## 2. Materials and Methods

### 2.1. Bacterial strains and cell lines

EPEC E2348/69 strains were grown in Luria Bertani (LB) (Sigma-Aldrich, St. Louis, Missouri, United States of America) broth or agar. Overnight bacterial cultures were grown at 37°C, 180 rpm shaking (liquid) or static (agar) and primed in Dulbecco’s modified eagle medium (DMEM) as described below for infections.

MIA PACA2 cells were maintained in DMEM (Sigma) with 10% (v/v) heat-inactivated fetal bovine serum (FBS) (Sigma) at 37°C with 5% CO_2_.

### 2.2. Infection of PDAC cells

EPEC infection was performed as described before in Zhong *et al*., 2020 (Zhong et al., 2020) with modifications. Briefly, EPEC strains were primed by diluting the overnight cultures 50× in non-supplemented DMEM (low glucose) and growing for 3 h static at 37°C with 5% CO_2_ to induce T3SS expression. Isopropyl β-d-1-thiogalactopyranoside (IPTG) (Sigma) at 1 mM was added to the bacterial culture 30 min before infection to induce effector gene expression from pSA10 plasmids.

Infection was carried out at a multiplicity-of-infection (MOI) of 50:1 in DMEM with 10% FBS. Infected cells were centrifuged at 700 g for 10 min and incubated for 2 h static at 37°C, 5% CO_2_. At 2 h post-infection, 200 μg/ml gentamicin (Sigma) or 250 μg/ml kanamycin (Sigma) (for EPEC-1-Tir_AA_ strain) was added to prevent bacterial overgrowth. Most further treatments and analysis was performed at or after 4 h post-infection, a timepoint previously shown to be sufficient for NleC expressed from both EPEC-0 and EPEC-1 background to exert similarly effective RelA depletion activities (Cepeda-Molero et al., 2017).

### 2.3. Cytokine and drug treatment

10 ng/ml TNFα (R&D Systems, Minneapolis, Minnesota, USA) was added to the cells 4 h post-infection (or at the equivalent timepoint in uninfected cells). 50 μM z-VAD-fmk (zVAD) (R&D Systems) and 5 μM necrosulfonamide (NSA) (Tocris) were added to the cells 30 min before infection.

For caspase-8 apoptosis positive control, 10 ng/ml TNFα (R&D Systems) and 10 μg cycloheximide (CHX) (Sigma) were added to the cells. For caspase-4 pyroptosis positive control, Ultrapure *E. coli* O111:B4 LPS (Invivogen, San Diego, California, USA) transfection was performed using Lipofectamine 2000 (Invitrogen, Waltham, Massachusetts, USA) at 5 μg/ml.

### 2.4. siRNA transfection

A total of 2.5×10^3^ cells/well were seeded in black clear-bottom 96-well plates 3 days prior to infection. siRNA (Dharmacon, Lafayette, Colorado, USA) mixed with Lipofectamine RNAiMAX (Invitrogen) in Opti-MEM were added to the cells 2 days before infection according to manufacturer’s instruction, with a final siRNA concentration of 20 nM. The medium was replaced with fresh DMEM with 10% FBS 1 day before infection.

### 2.5. Infection confirmation

At 2 h post-infection, infected cells were fixed by 4% paraformaldehyde for 15 min, washed by 3 PBS, permeabilised by 0.2% Triton X-100 (Sigma) for 4 min, washed again and blocked with 1% bovine serum albumin (BSA) (Sigma) for 10 min before being incubated with 1:1000 4’,6-diamidino-2-phenylindole (DAPI) (Sigma) and 1:200 Phalloidin Alexa-647 (Stratech, Ely, UK) for 30 min. Cells were then imaged using Opera Phenix High-Content Screening System (Perkin-Elmer, Waltham, Massachusetts, USA) at 63×.

To quantify infection rate, at 2 h post-infection, infected cells were washed by 3×PBS. Cells were lysed with Triton X-100 at 0.1% (non-lethal to bacteria), serially diluted and then plated on LB agar plates. Colony forming units (CFUs) were counted after overnight incubation at 37°C.

### 2.6. High content image acquisition and analysis

A total of 5×10^3^ cells/well were seeded in black clear-bottom 96-well plates 1 day prior to infection. Infection was performed as described above. High content imaging was performed on live cells using Opera Phenix High-Content Screening System (Perkin-Elmer) at 20× with 80% humidity, 37°C and 5% CO_2_. 4 fields per well and 3 wells per condition was imaged from 4 h post-infection (before TNFα addition) to 20 h post-infection with 15 min intervals. Image analysis was performed on Harmony software (Perkin-Elmer). Workflow was summarised in Figure S1. Nucleus and cytoplasm were segmented using PCNA-mScarlet fluorescence and RelA-eGFP fluorescence, respectively. Cells touching the image borders were removed from analysis. The remaining cells were filtered according to the presence of morphologies that correspond to cell death feature, including low cell size (cell shrinkage), high cell roundness (cell rounding), low nuclear size (nuclear shrinkage) and high nuclear/cytosolic area ratio (loss of cytoplasm volume). Dead cells were removed from analysis. The number of dead cells were generally not used for cell death rate calculation due to an increase in inaccuracy following the disintegration of cell debris overtime. The number of live cells were instead used for calculating percentage live cells compared to the 1^st^ timepoint as a readout of cell loss. Live cells were then used for measuring RelA nuclear intensity and nuclear/cytosolic ratio: nuclear region for intensity measurement was reduced by 2-pixel inward from the nucleus border; cytosolic intensity was measured in a ring region from 2-pixel to 6-pixel outward of the nuclear border.

Single cell tracking was performed using the LIM Tracker plugin (Aragaki et al., 2022) in ImageJ on at least 100 cells in total, exact number of cells were stated in each relevant figure. Tracking was terminated if a cell shows the abovementioned cell death morphologies in the next timepoint (and remains a dead cell for all following timepoints), if a cell undergoes division or if a cell reaches the imaging endpoint without the previous two events. If a cell moves out of the image border at any timepoint, it will not be used for analysis. Nuclear RelA measurements from 4 h to 6 h post-infection were used for analysis while the entire 20 h time course was used for cell fate determination.

### 2.7. LDH release assay

A total of 1×10^4^ cells/well were seeded in clear-bottom 96-well plates 1 day prior to infection. Infection was performed as described above. At 20 h post-infection, lactate dehydrogenase (LDH) release assay was performed using CytoTox 96 Non-Radioactive Cytotoxicity Assay kit (Promega, Madison, Wisconsin, USA) according to manufacturer’s instruction. Briefly, spent medium was mixed with CytoTox 96 Reagent at 1:1 ratio and incubated at room temperature for 30 min followed by 1 Stop Solution addition. Absorbance at 490 nm was measured and first normalised to the uninfected/untreated cells. Increase in percentage LDH release compared to uninfected was calculated by dividing each normalised reading over normalised positive control reading (cells treated with 1% Triton X-100).

### 2.8. Statistical analysis

All experiments were independently repeated at least 3 times as indicated in the figure legends. Each condition has 3 wells serving as technical repeats for each high content imaging experiment. Each condition has 2 wells serving as technical repeats for each LDH assay. Statistical analysis was performed using GraphPad Prism 5.1. 1-way or 2-way ANOVA followed by Tukey post-test or Bonferroni post-test, respectively, were performed on the means as listed in the figure legends. Significant result was defined as having a p-value of <0.05.

## 3. Results

### 3.1. EPEC effectors differentially regulate TNF_α_-mediated NF-_κ_B signalling dynamics

We first infected the model PDAC cell line MIA PACA2 with EPEC WT and EPEC-0 (mutant with all effector genes deleted (Cepeda-Molero et al., 2017)) for 2 h. Immunofluorescence showed actin polymerisation, indicative of actin pedestal formation, adjacent to the EPEC WT bacteria but not EPEC-0 (Figure 1A). Of the 275 EPEC WT-infected cells counted in total, we have only found 3 cells with no pedestal, with an average 99% pedestal formation rate. This confirms the successful EPEC infection and T3SS activity on MIA PACA2 cells.

**Figure 1.**
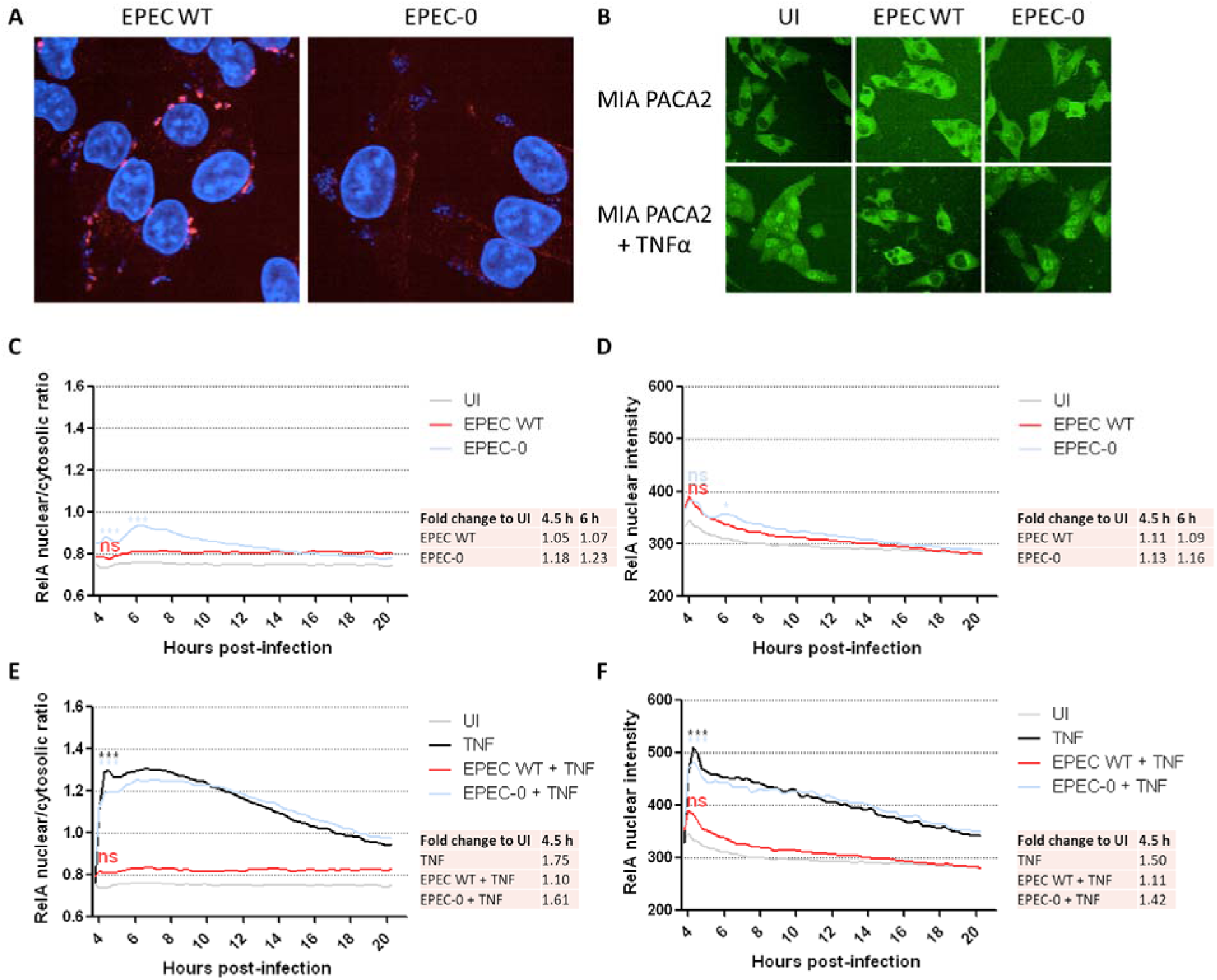
RelA nuclear/cytosolic ratio and nuclear intensity after wild-type EPEC infection on PDAC cells. (A) Immunofluorescence microscopy of MIA PACA2 cells infected by EPEC WT and EPEC-0 for 2 h. Blue: DAPI. Red: Phalloidin. Representative images from n=2 independent biological repeats were shown. (B) Live imaging of MIA PACA2 cells infected by EPEC WT and EPEC-0 or uninfected (UI). Infection was performed for 2 h followed by gentamicin treatment. TNF_α_ treatment was performed at 4 h post-infection. Representative images at 5 h post-infection from n=3 independent biological repeats were shown. (C-F) RelA nuclear/cytosolic ratio (C, E) and RelA nuclear intensity (D, F) of MIA PACA2 cells infected by EPEC WT and EPEC-0, with (E, F) or without (C, D) TNFα treatment at 4 h post-infection, were measured from 4 h to 20 h post-infection with 15 min interval. Average fold change compared to UI at specified timepoints was shown in table. Mean from n=3 independent biological repeats were shown. Statistical significance was determined using 2-way ANOVA with Bonferroni post-test. Statistical significance of the 4.5 h (C-F) and 6 h (C, D) timepoints compared to UI is shown. ***p≤0.001; ns: non-significant.

MIA PACA2 cell expressing eGFP-tagged RelA from the *RELA* locus and mScarlet-tagged PCNA (Butera et al., 2022) was used to monitor NF-κB dynamics following infection by effector mutants using live cell timelapse imaging. Workflow of infection experiments and imaging analysis are shown in Figure S1. Both the RelA nuclear/cytosolic ratio and RelA nuclear intensity were used as readouts of NF-κB activation. Infection with EPEC WT resulted in no significant increase in RelA nuclear/cytosolic ratio and peak RelA nuclear intensity arbitrary units (AU) in the infected cells (∼0.8 and 370 AU, respectively) compared to the uninfected untreated cells (UI) (∼0.7 and 330 AU, respectively) (Figure 1B-F). TNFα addition to the uninfected cells resulted in robust translocation of NF-κB into the nucleus with a peak nuclear/cytosolic ratio of 1.3 and peak nuclear intensity of 500 AU, which are roughly 1.75-fold and 1.5-fold higher than UI, respectively. In comparison, TNFα addition 4 h post-infection did not further promote RelA nuclear/cytosolic ratio or peak RelA nuclear intensity in EPEC WT-infected cells, which remained at around 1.1-fold compared to UI (Figure 1E, F). Thus, EPEC infection is an effective suppressor of NF-κB nuclear translocation following TNFα stimulation.

To validate that that the effects of EPEC on NF-κB translocation were due to injected effectors, we also infected cells with EPEC-0. EPEC-0 infection itself resulted in a low-level increase in RelA nuclear/cytosolic ratio to a peak of 0.93 at 6 h on its own, equivalent to 1.23-fold compared to UI, and could not affect TNFα-induced increase in RelA nuclear/cytosolic ratio and RelA nuclear intensity (Figure 1B-F). Thus, the effector cocktail injected by the EPEC WT is highly effective at suppressing NF-κB activation.

We then tested the roles of the individual effectors, NleC, NleE and Tir (Figure 2A), in blocking NF-κB translocation, by infecting cells with mutant EPEC strains that only express these single effectors (Figure 2B). Enumeration of colony forming units (CFU) of bacteria attachment to cells showed similar results between the strains expressing the three effectors individually (Figure S2). Overexpression of the NF-κB protease NleC from EPEC-0 (EPEC-0-NleC) resulted in reduction of RelA signal from the entire cell (Figure 2B), confirming previously reported data (Baruch et al., 2010; Yen et al., 2010). Although the RelA nuclear/cytosolic ratio in the EPEC-0-NleC-infected cells appeared elevated compared to UI (Figure 2C), NleC reduced both RelA nuclear intensity and cytosolic intensity to 50∼100 AU below the UI baseline most prominently after 5 h post-infection (Figure 2D, S3A, B). TNFα addition weakly elevated RelA nuclear/cytosolic ratio in EPEC-0-NleC-infected cells to 1.29-fold compared to UI, followed by a slow increase from 6 h to 14 h post-infection (Figure 2E). However, regardless of TNFα stimulation, NleC kept both the RelA nuclear intensity from 5 h to 12 h post-infection and the RelA cytosolic intensity throughout 20 h at least 50 AU below the baseline (Figure 2F, S3A, B). Thus, NleC depletes RelA from the infected cells to prevent NF-κB activation in both short- and long-terms.

**Figure 2.**
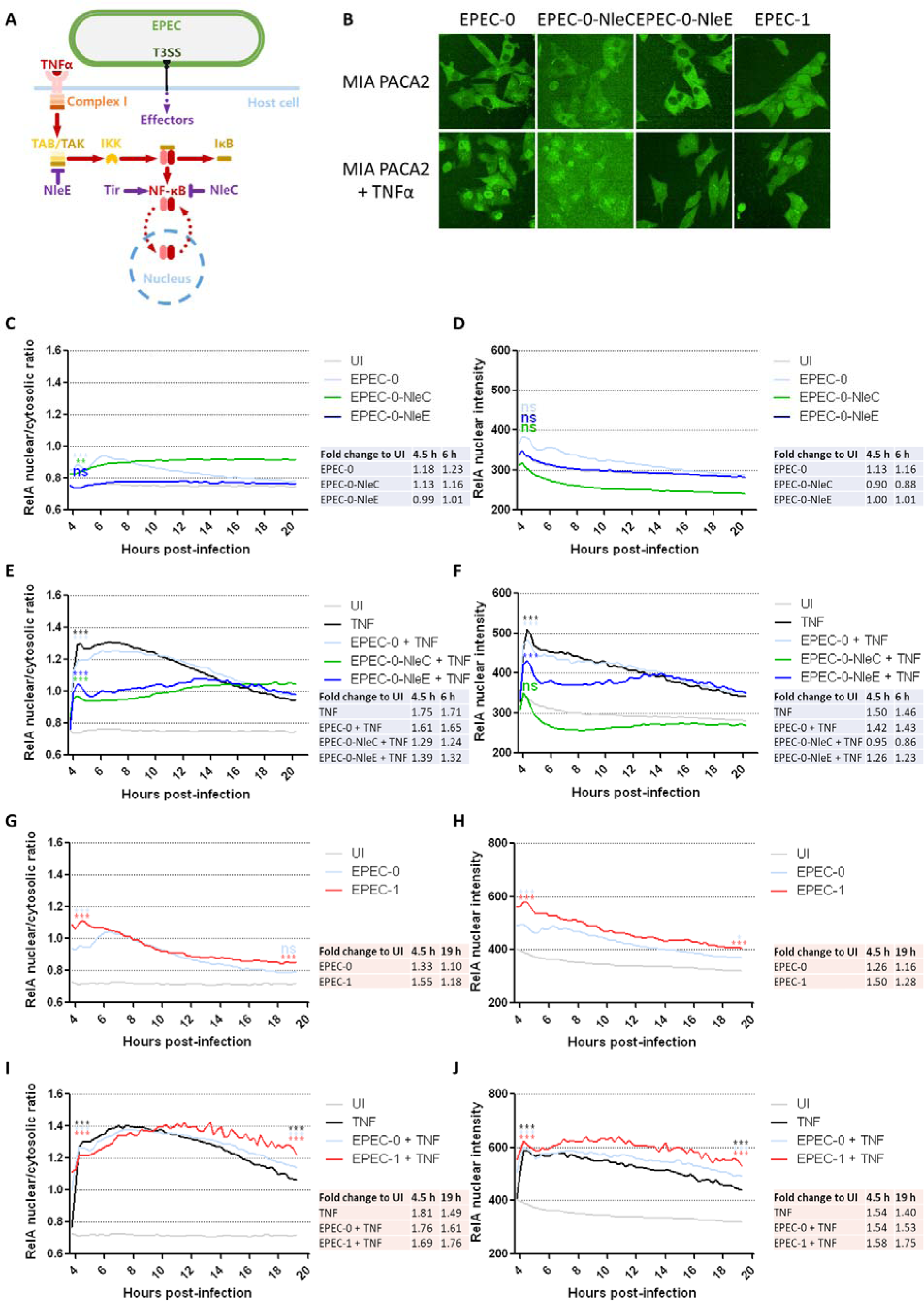
RelA nuclear/cytosolic ratio and nuclear intensity after EPEC single effector-expressing mutant infection. (A) Illustration of EPEC infection and known activities of NleC, NleE and Tir on TNFα-NF-κB signalling pathway. (B) Live imaging of MIA PACA2 cells infected by EPEC-0, EPEC-0-NleC, EPEC-0-NleE and EPEC-1 or uninfected (UI). Infection was performed for 2 h followed by gentamicin treatment. TNFα treatment was performed at 4 h post-infection. Representative images at 5 h post-infection from n=3 independent biological repeats were shown. (C-J) RelA nuclear/cytosolic ratio (C, E, G, I) and RelA nuclear intensity (D, F, H, J) of MIA PACA2 cells infected by EPEC EPEC-0, EPEC-0-NleC (C-F), EPEC-0-NleE (C-F) and EPEC-1 (G-J), with (E, F, I, J) or without (C, D, G, H) TNFα treatment at 4 h post-infection, were measured from 4 h to 20 h post-infection with 15 min interval. Average fold change compared to UI at specified timepoints was shown in table. Mean from n=3 independent biological repeats were shown. Statistical significance was determined using 2-way ANOVA with Bonferroni post-test. Statistical significance of the 4.5 h (C-J) and 19 h (G-J) timepoints compared to UI is shown. ***p≤0.001; **p≤0.01; ns: non-significant.

Overexpression of the TAB2/3 methyltransferase NleE from EPEC-0 (EPEC-0-NleE) resulted in a complete blockage of RelA nuclear/cytosolic ratio compared to EPEC-0 (Figure 2B, C), and its RelA nuclear intensity, as well as RelA cytosolic intensity, had no significant change compared to the uninfected untreated cells (Figure 2D, 32). After TNFα addition, the RelA nuclear/cytosolic ratio and RelA nuclear intensity in EPEC-0-NleE-infected cells have a reduced peak of 1.39-fold and 1.26-fold compared to UI followed by a slow increase from 6 h to 14 h post-infection (Figure 2E, F). Unlike NleC, NleE increased the RelA cytosolic intensity in the presence of TNFα in comparison to TNFα alone, most prominently from approximately 6 h to 10 h post-infection, corresponding to the same time the RelA nuclear intensity suppression was the strongest, confirming that NleE prevents the translocation process of RelA (Figure S3A, B). This confirms that NleE is another EPEC effector that blocks RelA activation in response to TNFα (Nadler et al., 2010; Newton et al., 2010; Zhang et al., 2012).

In comparison, we then tested how an EPEC NF-κB activator effector, Tir, modulates NF-κB activation in the absence or presence of TNFα. We infected MIA PACA2 cells with EPEC-1, with all effectors deleted except Tir (a.k.a. EPEC-0-Tir) (Cepeda-Molero et al., 2017) (Figure 2B). EPEC-1 promoted a moderate increase in RelA activation to around 1.5-fold compared to UI at 4.5 h post-infection (Figure 2G, H). Tir-dependent RelA activation is independent of its actin polymerisation ability, as the actin polymerisation-deficient mutant Tir Y474A/Y454A (Tir_AA_) has the same RelA nuclear/cytosolic ratio curve as the WT Tir (Figure S4A, B). In TNFα-treated cells, EPEC-1 was unable to significantly increase the peak RelA activation compared to EPEC-0, possibly suggesting that the RelA translocation after TNFα treatment has reached a maximum level. However, EPEC-1+TNFα promoted a more persistent nuclear RelA intensity after 6-8 h post-infection, which remained at 510 AU when close to infection endpoint, equivalent to 1.75-fold compared to UI, in contrast to 1.53-fold with EPEC-0+TNFα and 1.40-fold with TNFα alone (Figure 2I, J). We have also observed that the RelA cytosolic intensity in EPEC-1-infected TNFα-treated cells was 40 AU lower than UI but 40 AU higher than EPEC-0-infected TNFα-treated cells from 6 h to 12 h post-infection (Figure S4C, D). As an increase in both cytosolic and nuclear RelA intensity was observed, it suggests that Tir promotes a sustained production of RelA. Overall, Tir promotes NF-κB activation via a combination of NF- κB translocation and production; in the presence of TNFα, this leads to a more sustained NF-κB nuclear retention.

### 3.2. EPEC effectors synergise with TNF_α_ to promote cell death

TNFα can stimulate cell death pathways, thus we simultaneously quantified cell survival in parallel with NF-κB translocation following EPC infection. To quantify cell death in an automated fashion we devised a cell death morphology classifier, which measured cell shrinkage, cell rounding and nuclear condensation (Figure S1), to determine the percentage of cells with cell death morphologies at specific timepoints (Figure 3A, B). Using this classifier to exclude dead cells, live cells were counted and normalised to the 1^st^ timepoint as a readout of cell survival (Figure 3C, D). We first monitored cell survival during EPEC WT infection and TNFα treatment. No significant increase in cells with cell death morphologies was detected with EPEC WT infection; however, live cell percentages with EPEC WT and EPEC-0 infection were mildly lower by around 20% compared to uninfected untreated cells, suggesting a possible mild inhibitory effect of EPEC on cell growth (Figure 3A, C). With TNFα treatment alone, live cell percentage was reduced to around 80% (Figure 3B, D). Notably, EPEC WT, but not EPEC-0, blocked TNFα-induced cell death (Figure 3D). Thus, the full complement of EPEC effectors makes the host cell unable to respond to both TNFα-mediated activation of NF-κB (Figure 1) and cell death (Figure 3).

**Figure 3.**
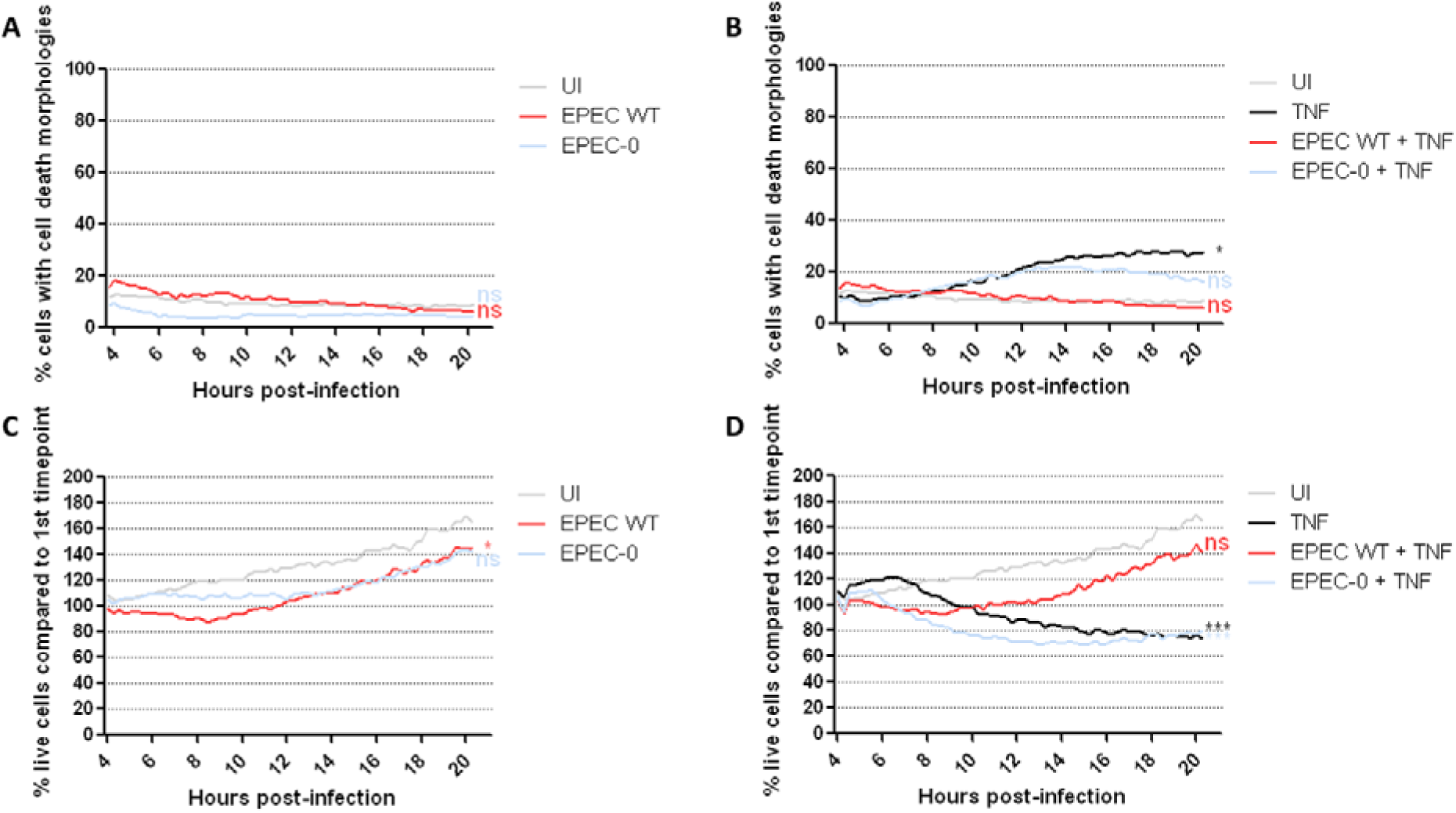
Changes in cell death morphology and cell survival after wild-type EPEC infection. Percentage cells with cell death morphologies (A, B) and Percentage live cells normalised to 1^st^ timepoint (C, D) of MIA PACA2 cells infected by EPEC WT and EPEC-0, with (B, D) or without (A, C) TNF_α_ treatment at 4 h post-infection, were measured from 4 h to 20 h post-infection with 15 min interval. Mean from n=3 independent biological repeats were shown. Statistical significance was determined using 2-way ANOVA with Bonferroni post-test. Statistical significance of the imaging endpoint compared to UI is shown. *p 0.05; ***p 0.001; ns: non-significant.

EPEC-0-NleC and EPEC-0-NleE reduce nuclear RelA in response to TNFα (Figure 1). We next tested if EPEC-0-NleC and EPEC-0-NleE are also involved in regulating cell survival downstream of TNFα. EPEC-0-NleC and EPEC-0-NleE did not change cell survival on their own, but synergised with TNFα, reducing the live cell percentage to 20% (Figure 4A, B). EPEC-1, on its own, appeared to be mildly toxic, which led to the reduction of live cell percentage to 60%. Similar to NleC and NleE, in TNFα-treated cells, EPEC-1 greatly reduced live cell number to 20% (Figure 4C, D). Although both Tir and TNFα alone has mild cytotoxicity, the combinatorial effect of Tir and TNFα on cell death occurred rapidly before 8 h post-infection, which was before the start of TNFα-induced cell death, suggesting a synergistic activity rather than a simply additive outcome.

**Figure 4.**
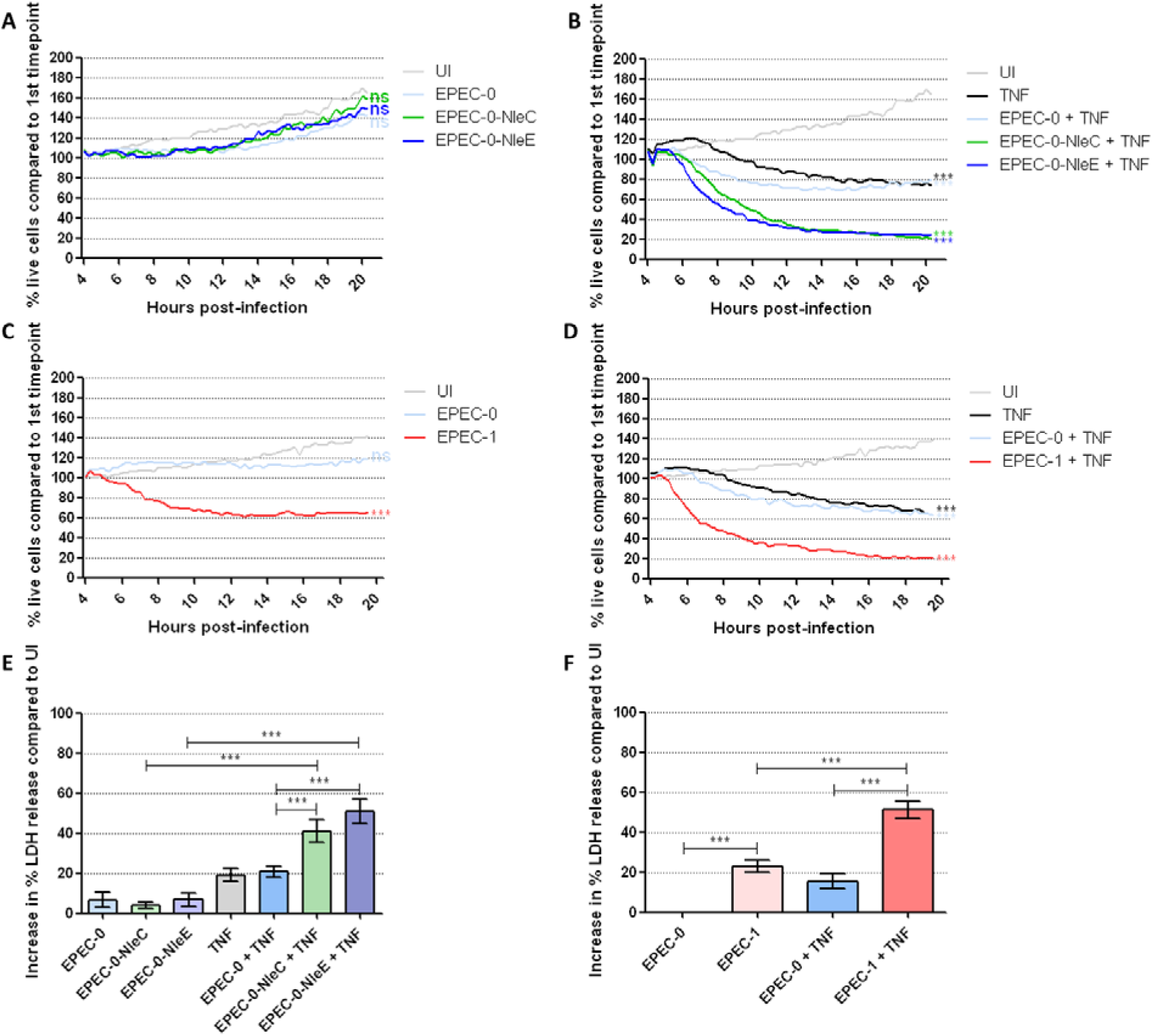
Cell survival and LDH release after EPEC mutant infection. (A-D) Percentage live cells (normalised to live cell numbers at 1^st^ timepoint) of MIA PACA2 cells infected by EPEC-0, EPEC-0-NleC (A, B), EPEC-0-NleE (A, B) and EPEC-1 (C, D), with (B, D) or without (A, C) TNF_α_ treatment at 4 h post-infection, were measured from 4 h to 20 h post-infection with 15 min interval. Mean from n=3 (A, B, C) and 4 (D) independent biological repeats were shown. Statistical significance was determined using 2-way ANOVA with Bonferroni post-test. Statistical significance of the imaging endpoint compared to UI is shown. (E, F) LDH release was measured in the supernatant of MIA PACA2 cells infected by EPEC-0, EPEC-0-NleC (E), EPEC-0-NleE (E) and EPEC-1 (F), with or without TNFα treatment at 4 h post-infection, at 20 h post-infection. Data were normalised to the uninfected untreated cells (as 0%) and Triton X-100-treated cells (as 100%). Means SEM from n=6 (E) and 8 (F) independent biological repeats are shown. Statistical significance was determined using 1-way ANOVA with Tukey post-test. ***p≤0.001; *p≤0.05; ns: non-significant.

Percentages of cells with cell death morphologies were not plotted for these samples with high cell death levels, due to the gradual disintegration of a large proportion of cell debris at later stages of infection which prevented their recognition. Instead, cell death levels were further validated using an endpoint LDH release assay at 20 h post-infection, which detects cell lysis events, including both late-stage apoptosis and necrosis. LDH assay confirmed that on their own, EPEC-0-NleC and EPEC-0-NleE have no toxicity, while EPEC-1 induced a mild increase on cell death; all three strains promoted LDH release to approximately 40-50% in the presence of TNFα (Figure 4E, F). In addition, Tir_AA_ induced moderately higher cell death compared to the wild-type Tir (Figure S4C-E). However, given that RelA translocation is unaffected by Tir_AA_ mutations, the increase in cell death with Tir_AA_ is likely unrelated to NF-κB.

Therefore, NleC, NleE and Tir, despite their opposite effects on NF-κB activation, all synergise with TNFα in causing cell death.

### 3.3. Caspase-8 is universally involved in cell death induced by all three effector-TNF_α_ combinations

To identify the cell death mechanism induced by the effector-TNFα combinations, we first used pan-caspase inhibitor z-VAD-fmk (zVAD) and necroptosis protein MLKL inhibitor necrosulfonamide (NSA) to investigate the caspase-dependence of these events. Treatment by zVAD, but not NSA, completely blocked cell death in EPEC-0-NleC- and EPEC-0-NleE-infected TNFα- treated cells (Figure 5A). Similarly, zVAD but not NSA significantly reduced cell death in EPEC-1-infected cells with or without TNFα treatment (Figure 5B), thus NleC, NleE, and Tir all promote caspase-dependent cell death.

**Figure 5.**
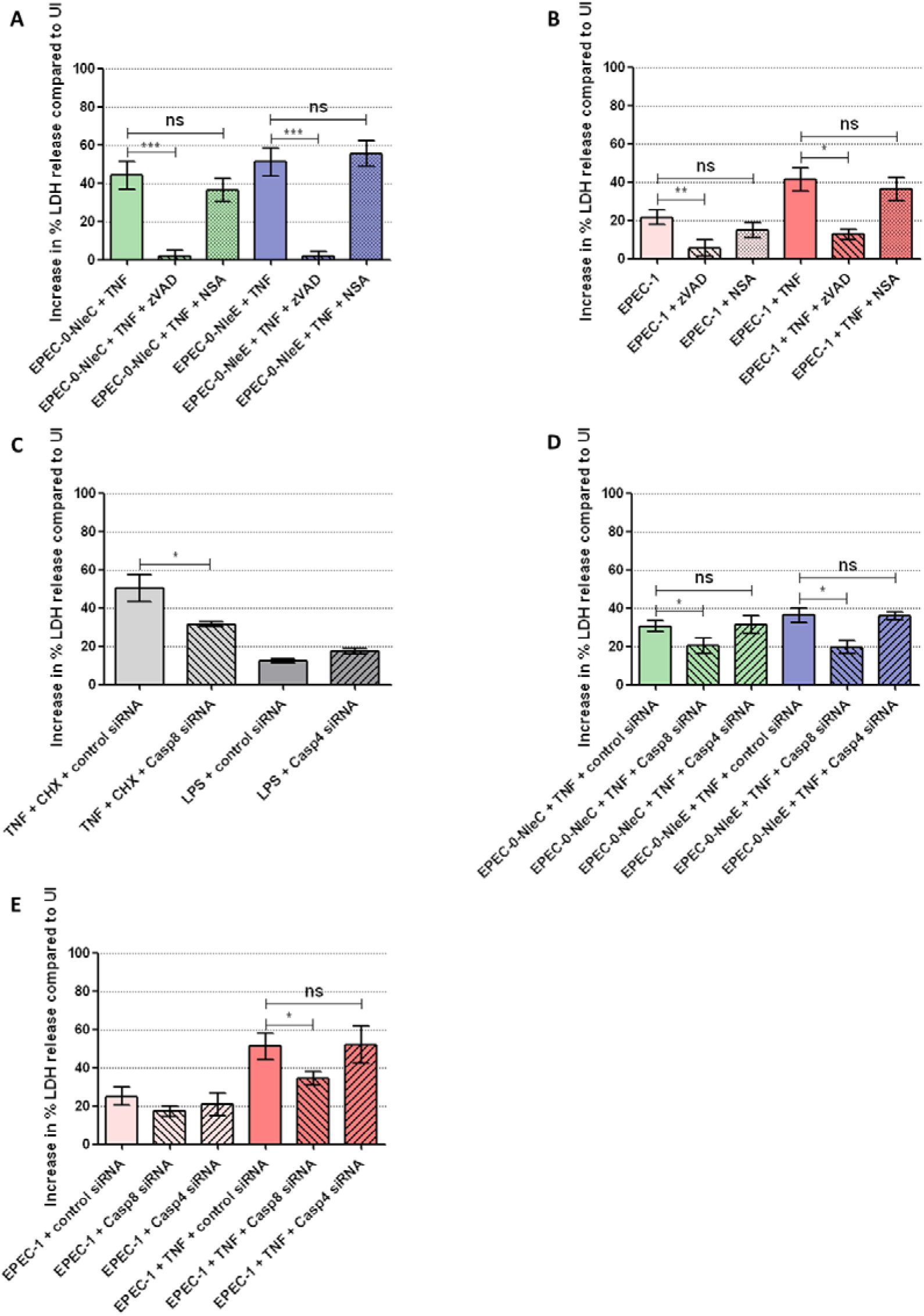
Changes in effector-TNF_α_-dependent cell death after caspase inhibitor and siRNA treatment. (A, B) LDH release was measured in the supernatant of MIA PACA2 cells infected by EPEC-0-NleC and EPEC-0-NleE with TNFα treatment (A), or EPEC-1 with or without TNFα treatment (B). zVAD and NSA treatment was performed 30 min prior to infection. Means SEM from n=4 independent biological repeats are shown. (C) MIA PACA2 cells were transfected by control, caspase-8 or caspase-4 siRNA 3 days before infection. LDH release was measured in the supernatant of siRNA-transfected MIA PACA2 cells treated by TNFα and CHX for 20 h or LPS lipofectamine transfected for 20 h. Means SEM from n=4 independent biological repeats are shown. (D, E) LDH release was measured in the supernatant of control, caspase-8- or caspase-4-silenced MIA PACA2 cells infected by EPEC-0-NleC and EPEC-0-NleE with TNFα treatment (D), or EPEC-1 with or without TNFα treatment (E). Means SEM from n=5 (D) and 4 (E) independent biological repeats are shown. Statistical significance was determined using 1-way ANOVA with Tukey post-test. *p≤0.05; **p≤0.01; ***p≤0.001; ns: non-significant.

Death receptor signalling can activate caspase-8, which can lead to extrinsic apoptosis or pyroptosis, and caspase-10, separately leading to extrinsic apoptosis (Wang et al., 2001, 2008). Meanwhile, NF-κB promotes caspase-4 expression (Yang et al., 2015), which participates in non-canonical inflammasome formation and pyroptosis. Among the initiator caspases, caspase-4, caspase-8 and caspase-10 are detectable in MIA PACA2, while caspase-1 (canonical inflammasome), caspase-5 (non-canonical inflammasome) and caspase-9 (intrinsic apoptosis) are not detectable (Bernhard et al., 2022). Additionally, caspase-10 cannot be inhibited by zVAD (Lafont et al., 2010). Therefore, we chose caspase-8 and caspase-4 siRNA for further investigations. According to the positive controls, using TNFα and cycloheximide (CHX) co-treatment to activate caspase-8, caspase-8 siRNA silencing significantly reduced TNFα/CHX-dependent apoptosis; while LPS transfection, which can activate rapid caspase-4-dependent pyroptosis, had little effect on cell death showing only 10% LDH increase even after 20 h, suggesting that the caspase-4 signalling pathway is deficient in this cell line (Figure 5C). Consistent with the positive controls, caspase-8 silencing reduced cell death caused by EPEC-0-NleC+TNFα and EPEC-0-NleE+TNFα as well as EPEC-1+/-TNFα treatment, while caspase-4 silencing remained ineffective (Figure 5D, E). Therefore, NleC, NleE and Tir can all promote caspase-8-mediated cell death in PDAC cell lines treated by TNFα, even though they differentially regulate NF-κB translocation following the same stimulation.

### 3.4. Single cell RelA nuclear intensity tracking reveals sub-populational variations in response to EPEC infection

To further analyse the correlation between effector-dependent NF-κB regulation and cell death within a population, we tracked single cells infected by EPEC-0-NleC, EPEC-0-NleE or EPEC-1 and treated by TNFα. We only quantified RelA and death in cells directly exposed to live EPEC, which we term 1^st^ generation cells, as later generations only partially inherit the effectors from their parent cells. Cells were tracked from 4 h (with TNFα addition immediately after the 4 h timepoint) till 20 h post-infection, cell division or cell death. 1^st^ generation cells were classified into surviving and dead sub-populations. Surviving cells included those that underwent cell division, while dead cells were those that underwent cell shape changes characteristic of cell death, including cell shrinkage, cell rounding and nuclear condensation at any point during the time course. We also tracked the nuclear RelA intensity in the surviving and dead sub-populations before 6 h post-infection, when most 1^st^ generation cells are still alive to track.

Single cell tracks were normalised to the average intensity of the first image to minimise exposure differences between experiments. Overall, single cell tracks show a large variation in RelA intensity, ranging from ∼50 AU (50% of the average intensity) to ∼200 AU (200% of the average intensity) (Figure 6). Cells that survived EPEC-0-NleC+TNFα co-treatment had significantly higher overall RelA nuclear intensity compared to the dead sub-population at the time of, and throughout, the time course (Figure 6A, B, S5). A small number of cells with persistent RelA activation, with around 140 AU normalised RelA nuclear intensity, appeared to be completely resistant to NleC-dependent cell death, while cells with peak RelA nuclear intensity lower than 80 AU were all sensitive to NleC-dependent cell death (Figure 6A). These data suggest that NleC targets RelA for degradation to levels below a critical threshold, which could blunt RelA activation in both short-term (4 h) and long-term (4-6 h), leading to cell death. Indeed, survival versus death can be predicted on RelA levels alone prior to TNFα treatment.

**Figure 6.**
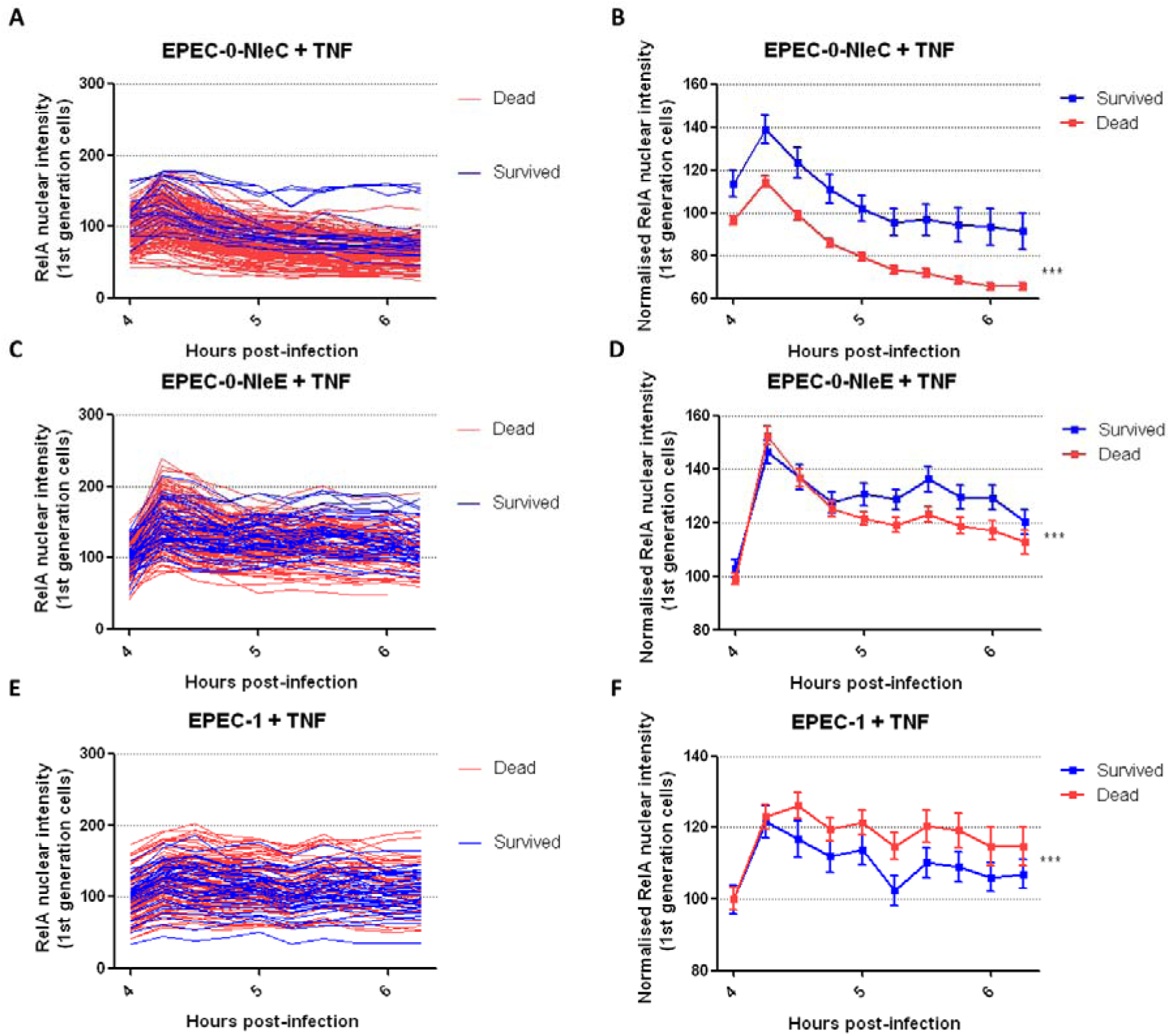
Single cell tracking of survived and dead populations after EPEC infection and TNF_α_ treatment. Single cell tracking data of 1^st^ generation MIA PACA2 cells infected by EPEC-0-NleC (A, B), EPEC-0-NleE (C, D) and EPEC-1 (E, F) with TNF_α_ treatment were classified based on cell fate: survived (including division) or dead before 20 h post-infection. RelA nuclear intensity data of each image were normalised to the average value of all cells at the 1^st^ timepoint to remove exposure differences between images. Single cell tracks are presented in A, C and D; Means SEM are presented in B, D, F. Numbers of cells tracked in total are 155 (20 survived and 135 dead) in EPEC-0-NleC+TNFα, 155 (40 survived and 115 dead) in EPEC-0-NleE+TNFα and 130 (40 survived and 90 dead) in EPEC-1+TNFα, each from 2 independent biological repeats. Statistical significance was determined using 2-way ANOVA. ***p×0.001.

Survived and dead sub-populations with EPEC-0-NleE+TNFα co-treatment did not differ in their average nuclear RelA intensity at the beginning of the tracking. In the short-term there was also no difference in RelA accumulation in cells that would ultimately die versus survive (4 h). However, after 45 min, the population bifurcated into surviving cells with high RelA levels, and dying cells with lower RelA levels, although the difference appears much less pronounced compared to EPEC-0-NleC+TNFα (Figure 6C, D, S6). Cells with persistent RelA activation, with over 160 AU normalised RelA nuclear intensity during 5-6 h post-infection, were also showing high resistance to NleE-dependent cell death (Figure 6C). Thus, we conclude NleE promotes cell death by suppressing the activation of RelA at a later timepoint compared to NleC. Following NleE treatment and RelA activation, there is a bifurcation point, leading to a sub-population that will die, and the other that will survive. This decision is not due to pre-existing differences in RelA, but rather could be due to intrinsic variations in feedback control on RelA or extrinsic variations in cell state (Prescott et al., 2021).

Similarly in the short-term, the effect of Tir on RelA was not different between cells that ultimately survived or died. Consistent with the idea that Tir enhances RelA-mediated cell death, we observed that following EPEC-1+TNFα co-treatment, the sub-population that survived had lower RelA intensity in the long-term (4-6 h post-infection) compared to the dead population (Figure 6E, F, S7). Cells with persistent RelA activation, with over 160 AU of normalised RelA nuclear intensity, were highly sensitive to Tir-dependent cell death (Figure 6E). This is opposite to what we have observed with EPEC-0-NleC+TNFα and EPEC-0-NleE+TNFα. Thus, in contrast to NleC and NleE, this supports the idea that Tir promotes cell death by directly or indirectly upregulating RelA.

### 3.5. NleC and NleE reduce Tir-dependent RelA activation

We observed that sub-populations of MIA PACA2 cells with higher RelA intensity after infection and TNFα treatment are less susceptible to NleC and NleE but more susceptible to Tir. Tir and NleC/NleE, in the presence of TNFα, likely induce cell death via different mechanisms through differential regulation of NF-κB. Thus, the use of single effectors allows us to ‘tune’ TNFα-NF-κB signalling in ways that either promote death by decreasing NF-κB (NleC and NleE) or by increasing NF-κB (Tir). We thus sought to determine if TNFα-NF-κB signalling could be re-engineered using combinations of effectors. We infected the PDAC cell lines with EPEC-1-NleC and EPEC-1-NleE, with or without TNFα (Figure 7A). CFU assay confirmed that the expression of NleC or NleE from EPEC-1 did not affect its ability to attach to cells (Figure S8). As expected, NleC reduced Tir-dependent increase in RelA activation and depleted RelA nuclear intensity (Figure 7B-E). In comparison, NleE also reduced Tir-dependent increase in RelA activation from 1.5-fold compared to UI to around 1.1-fold in the absence of TNFα (Figure 7B, C). In the presence of TNFα, RelA activation with EPEC-1-NleE+TNFα were reduced at the start of TNFα addition compared to EPEC-1+TNFα, from around 1.6-fold compared to UI to 1.29-fold, but gradually increasing from 6 h to 10 h post-infection to over 1.8-fold compared to UI, close to EPEC-1+TNFα (Figure 7D, E). Therefore, NleE blocks Tir- and TNFα-induced early peak RelA translocation 4-5 h post-infection but is less effective at blocking Tir- and TNFα-induced RelA translocation 6-10 h post-infection. Taken together, this suggests that the combination of Tir+NleE modulates NF-κB dynamics downstream of TNFα stimulation by inhibiting immediate translocation of RelA, but sustaining an increasing RelA translocation at later stage of infection.

**Figure 7.**
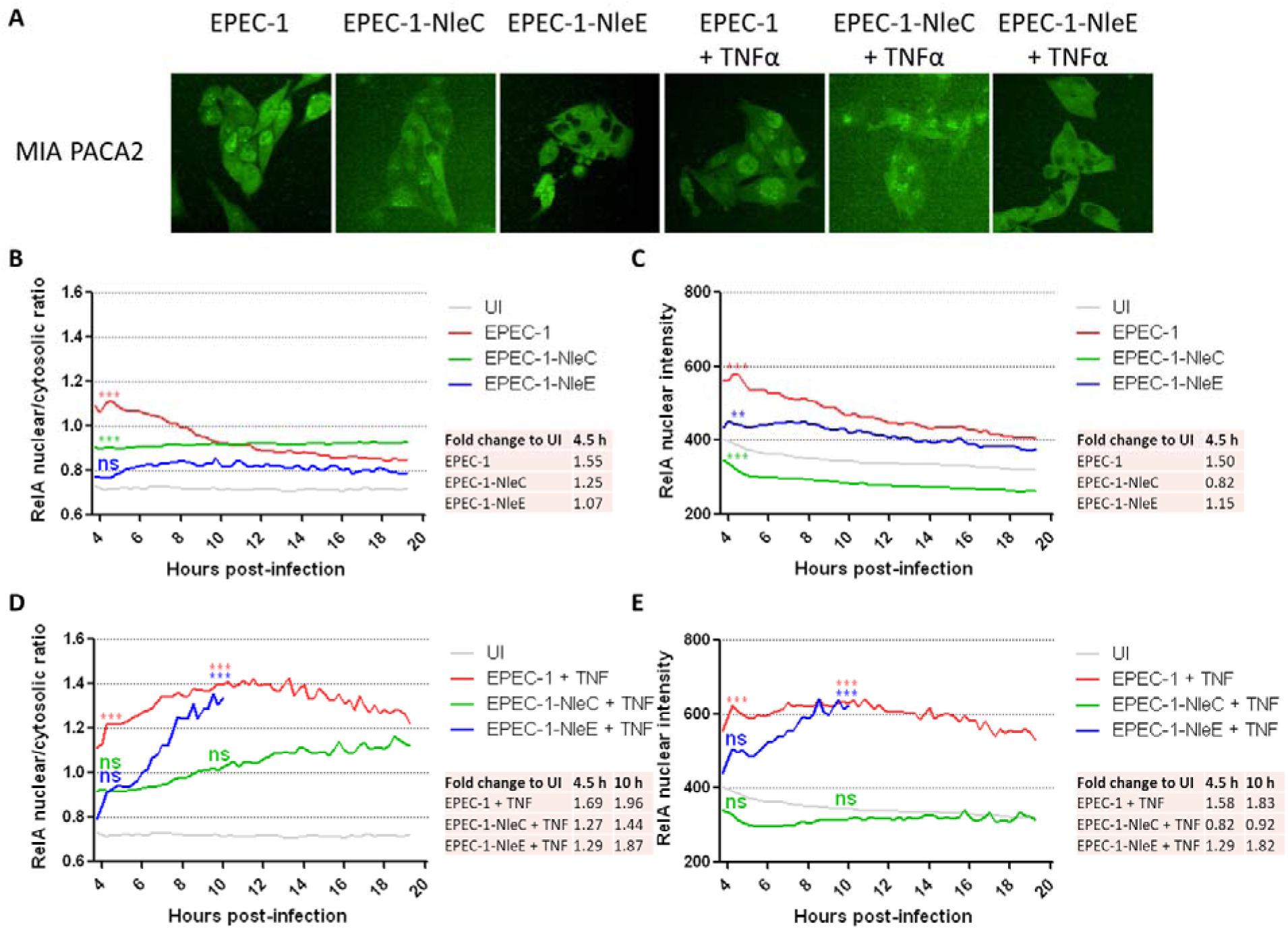
RelA nuclear/cytosolic ratio and nuclear intensity in cells with effector combinations. (A) Representative images at 5 h post-infection of MIA PACA2 cells infected by EPEC-1, EPEC-1-NleC and EPEC-1-NleE, with or without TNF_α_ treatment at 4 h post-infection, from n=3 independent biological repeats. (B-E) RelA nuclear/cytosolic ratio (B, D) and RelA nuclear intensity (C, E) of MIA PACA2 cells infected by EPEC-1, EPEC-1-NleC and EPEC-1-NleE, with (D, E) or without (B, C) TNF_α_ treatment at 4 h post-infection. Data for EPEC-1-NleE+TNF_α_ after 10 h post-infection were not plotted due to insufficient cell number. Average fold change compared to UI at specified timepoints was shown in table. Mean from n=3 independent biological repeats were shown. Statistical significance was determined using 2-way ANOVA with Bonferroni post-test. Statistical significance of the 4.5 h peak (B-E) and 10 h (D, E) timepoints compared to UI is shown. ***p×0.001; **p×0.01; ns: non-significant.

### 3.6. NleC and NleE differentially regulate Tir-dependent cell death

As NleC and NleE in combination with Tir reduce RelA activation with different dynamics, we then measured cell death in EPEC-1-NleC- and EPEC-1-NleE-infected cells. EPEC-1-NleC infection led to increased cell survival and decreased LDH release compared to EPEC-1 alone in the absence of TNFα (Figure 8A, G), showing that NleC antagonises Tir’s ability to promote cell death. Furthermore, EPEC-1-NleC and EPEC-1 infection resulted in similar levels of cell survival and LDH release in the presence of TNFα (Figure 8B, G). Single cell tracking revealed that the survived sub-population with EPEC-1-NleC+TNFα had higher RelA nuclear intensity compared to the dead sub-population (Figure 8C, D, S9), a similar trend as EPEC-0-NleC+TNFα. This indicates that the NleC-dependent RelA depletion prevented Tir’s ability to induce cell death which likely requires RelA activation. Therefore, Tir and NleC has no synergistic effects on cell death. As NleC depletes RelA, this also confirms that Tir requires RelA for its cytotoxicity.

**Figure 8.**
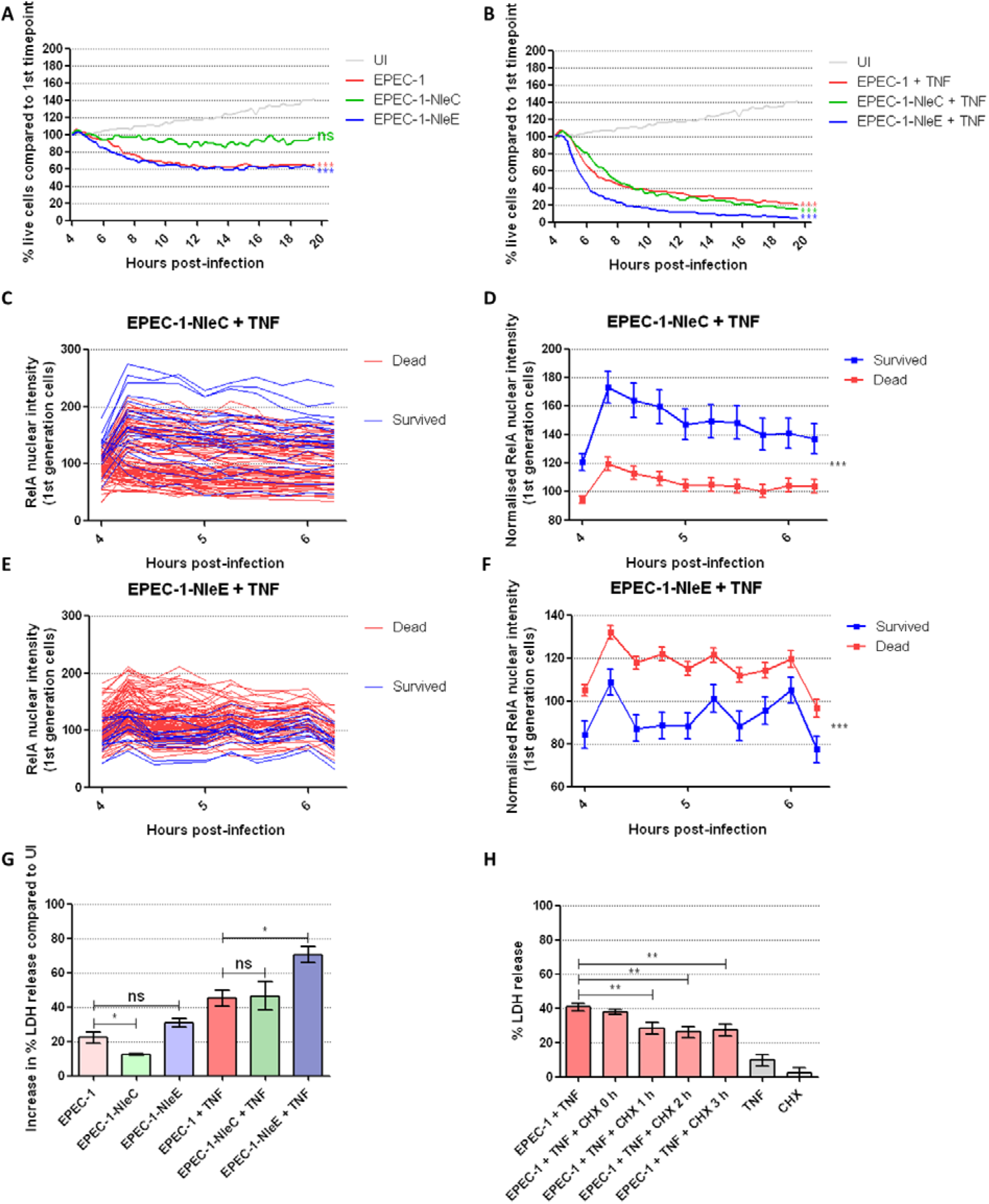
Cell death in cells with effector combinations. (A, B) Percentage live cells of MIA PACA2 cells infected by EPEC-1, EPEC-1-NleC and EPEC-1-NleE, with (B) or without (A) TNFα treatment at 4 h post-infection. Mean from n=3 independent biological repeats were shown. Statistical significance was determined using 2-way ANOVA with Bonferroni post-test. Statistical significance of the imaging endpoint compared to UI is shown. (C-F) Single cell tracking data of 1^st^ generation MIA PACA2 cells infected by EPEC-1-NleC (C, D) and EPEC-1-NleE (E, F) with TNFα treatment. Single cell tracks are presented in C and E; Means SEM are presented in D and F. Numbers of cells tracked in total are 110 (20 survived and 90 dead) in EPEC-1-NleC+TNFα and 125 (15 survived and 110 dead) in EPEC-1-NleE+TNFα from 2 independent biological repeats. Statistical significance was determined using 2-way ANOVA. (A) (G) LDH release was measured in the supernatant of MIA PACA2 cells infected by EPEC-1, EPEC-1-NleC and EPEC-1-NleE with or without TNFα treatment. Means ± SEM from n=4 independent biological repeats are shown. Statistical significance was determined using 1-way ANOVA. (B) (H) LDH release was measured in the supernatant of MIA PACA2 cells infected by EPEC-1 with TNFα treatment at 4 h post-infection. CHX treatment was performed either simultaneously as TNFα or at 1 h, 2 h or 3 h after TNFα treatment. Means SEM from n=4 independent biological repeats are shown. Statistical significance was determined using 1-way ANOVA. ***p≤0.001; **p≤0.01; *p≤0.05; ns: non-significant.

Although NleE had little effect on Tir-dependent cell death in the absence of TNFα (Figure 8A, G), in the presence of TNFα, NleE was synergistic with Tir in promoting death. Following EPEC-1-NleE+TNFα treatment, cell death occurred more rapidly compared to EPEC-1+TNFα with a 95% decrease in survival and 70-80% LDH release at 20 h post-infection (Figure 8B, G). The survived sub-population with EPEC-1-NleE+TNFα had lower RelA nuclear intensity compared to the dead sub-population (Figure 8E, F, S10), a similar trend as EPEC-1+TNFα, possibly suggesting that the cell death mechanism is mainly attributed to Tir activity. Furthermore, unlike the dead and survived sub-populations with EPEC-1+TNFα or EPEC-0-NleE+TNFα which bifurcate after the RelA peak, the differences between the two sub-populations with EPEC-1-NleE+TNFα occurred throughout 4-6 h post-infection (Figure 8E, F, S10). This difference indicates that the NleE’s ability to block pro-survival actions of RelA within the first 30 min to 1 h after TNFα treatment may be important to the increase in Tir-dependent cell death, while the long-term RelA translocation induced by Tir and TNFα and are not blocked by NleE leads to pro-cell death functions of RelA.

In addition, EPEC-0-NleC and EPEC-0-NleE co-infection did not increase cell death more than EPEC-0-NleC or EPEC-0-NleE alone, with or without TNFα (Figure S11), likely due to the fact that both effectors target the similar low RelA sub-populations according to previous single cell tracking results, or that the strong RelA depletion outcome of NleC negates any chance for NleE to exert additional inhibitory effect.

Gene expression patterns during NF-κB activation change overtime. Different set of RelA targets are induced in the immediate response to TNFα vs the long-term (Tian et al., 2005; Zhao et al., 2018). Therefore, we hypothesised that as Tir promotes RelA persistence, the increased late-stage RelA-dependent gene expression is responsible for the increase in TNFα-dependent cell death. Moreover, we proposed that NleE reduces the expression of the early-stage pro-survival genes, but not the late-stage Tir-induced RelA-dependent genes to further enhance cell death. We treated cells with EPEC-1+TNFα with the protein synthesis inhibitor CHX, either simultaneously with TNFα, or at 1-3 h after TNFα addition. Simultaneous TNFα and CHX treatment would block both early- and late-stage gene expression while delayed CHX treatment would only block the late-stage (1 h, 2 h or 3 h post-TNFα treatment) gene expression in EPEC-1+TNFα-treated cells. All delayed CHX addition after 1 h of incubation with TNFα resulted in only around 0.6∼0.7-fold LDH release compared to EPEC-1+TNFα with or without simultaneous CHX addition (Figure 8H). This supports the idea that EPEC-1+TNFα-induced gene expression within 1 h of TNFα treatment reduces cell death and/or promotes survival, while gene expression after 1 h of TNFα treatment likely promotes cell death. Therefore, it is possible for NleE-dependent early-stage RelA inhibition to reduce pro-survival gene expression, while enabling Tir-induced cell death, which relies on a persistent RelA nuclear retention, to proceed.

## 4. Discussion

In this study, we have used several NF-κB-regulating EPEC effector proteins, Tir, NleC and NleE, and their combinations to differentially regulate NF-κB activation and cell death downstream of the TNFα signalling pathway in PDAC cells. These effectors differentially regulate both NF-κB signalling and cell death: Tir activates NF- κB (Zhong et al., 2020), NleC degrades NF-κB (Yen et al., 2010) and NleE inhibits NF-κB nuclear translocation via TAB2/3 modification (Zhang et al., 2012); while Tir is moderately cytotoxic, NleC and NleE do not induce cell death on their own. TNFα treatment greatly enhanced cell death levels in combination with each of the three effectors, despite distinctive differences in their NF-κB regulation profiles. Tracking of the survived and dead sub-populations revealed that higher average RelA nuclear intensity is a trait of the survived populations of EPEC-0-NleC/NleE+TNFα, both of which reduce TNFα-dependent increase in nuclear RelA; oppositely, lower average RelA nuclear intensity is associated with the survived sub-population of EPEC-1+TNFα, which promotes persistence in nuclear RelA. NleC and NleE likely promotes cell death by preventing TNFα-dependent RelA activation whereas Tir-dependent cell death requires RelA activation (Figure 9).

**Figure 9.**
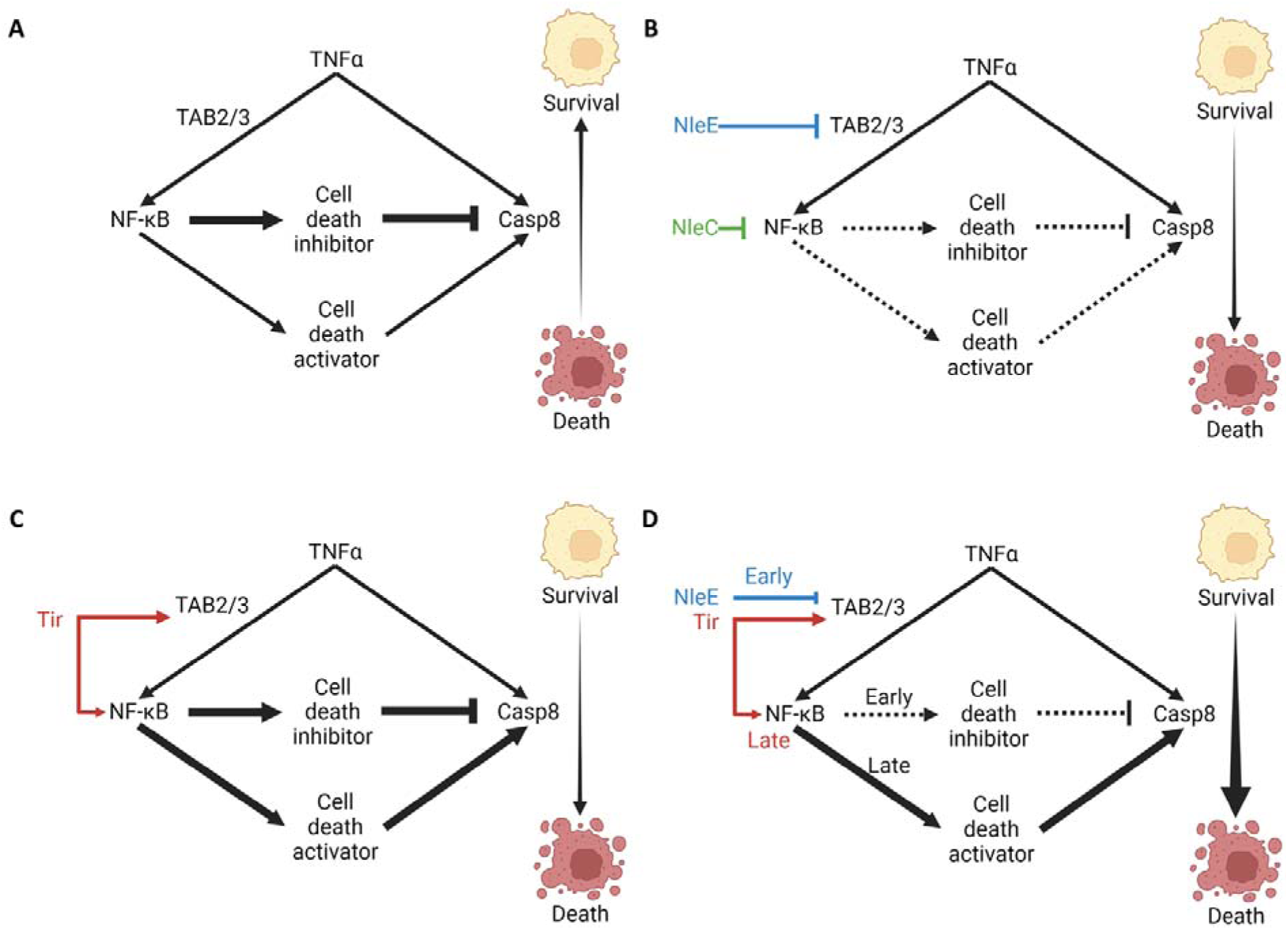
Model of NF-_κ_B and cell death regulation by effector-TNF_α_ combinations. (A) TNF_α_ signalling alone activates NF-_κ_B and recruits caspase-8, which is inhibited by NF-_κ_B-transcribed cell death inhibitors. Meanwhile, NF-_κ_B can also transcribe cell death activators. However, TNF_α_-treated cells only have low level of cell death, suggesting that the overall effect of the cell death inhibitors outweighs that of the cell death activators. (B) NleE inhibits TAB2/3 while NleC degrades NF-_κ_B, blocking NF-_κ_B-dependent transcription to free caspase-8 downstream of TNF_α_ signalling, leading to higher cell death. (C) Tir activates NF-κB via both TAB2/3-dependent NF-κB activation and increases cellular NF-κB production, leading to a more persistent NF-κB activation without a further increase in peak NF-κB activation in addition to TNFα signalling. It is likely that this disrupts the balance of NF-κB-transcribed cell death inhibitors and activators and the overall effect of the cell death activators now outweighs that of the cell death activators. (D) NleE inhibits TAB2/3, which leads to reduction of Tir- and TNFα-dependent peak NF-κB activation. As the NF-κB-transcribed cell death inhibitors tend to have transient expression, this enables caspase-8 activation to proceed. NleE-dependent NF-κB suppression is also transient, allowing a later increase in Tir-dependent NF-κB activation, further enhancing cell death. This figure was created with BioRender.com.

TNFα engages in both anti-apoptotic and pro-apoptotic signalling pathways. Upon stimulation, TNFα receptor recruits TRADD, TRAF2, RIPK1 and cIAP1/2 to form Complex I, and later TRADD dissociates to form Complex II with FADD and caspase-8 (Hsu et al., 1996; Micheau and Tschopp, 2003). These branches of TNFα signalling are tightly regulated. Caspase-8 inhibition by cFLIP and RIPK1 ubiquitination by cIAPs allow the recruitment of TAB and TAK, leading to NF-κB activation (Micheau et al., 2001; Moulin et al., 2012); while the loss of NF-κB-transcribed anti-apoptotic proteins allows caspase-8 activation (Hsu et al., 1996; Micheau and Tschopp, 2003; Wang et al., 2008). Meanwhile, NF-κB also transcribes cell death activators (Singh et al., 2007; Zheng et al., 2001) and overactivation of NF-κB can lead to apoptosis (Ricca et al., 2001; Ryan et al., 2000). Furthermore, different NF-κB target genes can be transcribed with different dynamics. Previous studies have identified pro-inflammatory cytokines including CXCL1, CXCL3 and IL-8, and anti-apoptotic proteins, such as A20, among the most rapidly transcribed within 1 h of NF-κB stimulation followed by a sharp reduction of their mRNA levels (Tian et al., 2005). In addition, the mRNA level of anti-apoptotic Bcl family protein Bcl-3 peaks at 1 h after NF-κB stimulation and remains unchanged before 3 h (Tian et al., 2005). Other anti-apoptotic proteins, such as cIAP1 and TRAF2, have continued transcription throughout 0 to 3 h post-stimulation before their reduction (Tian et al., 2005). In comparison, pro-apoptotic proteins, such as TRAF3, TRIM16 and IL-27 receptor, as well as adaptive immune response and antigen presentation proteins, have a delayed expression pattern, increasing only after 1 h of NF-κB stimulation and require more than 3 h to reach their maximal levels (Tian et al., 2005; Zhao et al., 2018). Therefore, a balance of cell death inhibitors and activators is required for cells under NF-κB stimulation to survive and the early induction of cell death inhibitors likely establishes an overall pro-survival profile for NF-κB (Figure 9A).

In our study, we believe that the NF-κB-regulating effectors disrupt this balance to switch the TNFα-stimulated cells from a low cell death state to a high cell death state. NleC and NleE likely promote TNFα-dependent cell death via preventing NF-κB-dependent transcription, leading to a reduction of caspase-8 inhibitors (Figure 9B). On the other hand, Tir promotes NF-κB activation on its own, likely through a TAB-dependent pathway similar as TNFα. Although Tir does not enhance the peak TNFα-induced RelA translocation, it promotes the persistence of nuclear RelA retention from 8 h to 20 h post-infection and induces RelA production in the nucleus from 6 h to 12 h post-infection. This delayed effect on RelA potentially favours the expression of genes involved in cell death activation. RelA is required for the cytotoxicity of Tir, as RelA depletion by NleC downregulates Tir-dependent cell death. It is possible that Tir elevates TNFα-dependent cell death via promoting a more persistent NF-κB activation, shifting the balance of NF-κB-transcribed pro-survival and cell death genes (Figure 9C).

Interestingly, we have previously seen that in colorectal cancer (CRC) cell line used as intestinal epithelial cell model, Tir induces Ca^2+^ and LPS influx to activate caspase-4, leading to GSDMD cleavage and pyroptosis with no caspase-8 involvement (Zhong et al., 2020, 2022). But in PDAC cells, caspase-8 plays a significant role in Tir-dependent cell death. LPS transfection, a classical method of caspase-4 activation, could not induce cell death in PDAC cells. This indicates that additional proteins essential for caspase-4 function may be missing in the PDAC cells. A possible candidate is the guanylate-binding proteins (GBPs), required for binding to LPS on transfected LPS aggregates or intracellular bacterial surface and promoting caspase-4 inflammasome assembly (Fisch et al., 2019, 2020; Pilla et al., 2014; Santos et al., 2018, 2020). Identifying the mechanism of resistance of PDAC cells to caspase-4-dependent cell death may potentially enable the effectors to obtain more cell death regulation functions.

Moreover, combining these effectors have surprising outcomes. The combination of NleC and NleE, both inhibitors of the NF-κB signalling pathway, has no synergistic or additive effect in cell death. It is likely that this is due to an overlap in their effects on RelA and their respective susceptible populations. On the other hand, EPEC-1-NleC-infected cells have depleted RelA nuclear intensity and reduced cell death compared to EPEC-1-infected cells. We believe that Tir-dependent RelA activation is essential for its cell death function on its own in PDAC cells, as NleC-dependent RelA degradation blocks the cytotoxicity of Tir. While NleC cannot synergise with Tir in cell death in the presence of TNFα, the RelA nuclear intensity difference between the survived and dead populations of EPEC-1-NleC+TNFα is similar to that of EPEC-0-NleC+TNFα while opposite to that in EPEC-1+TNFα, further suggesting that NleC antagonises Tir’s ability to promote cell death by preventing long-term RelA activation. In contrast, EPEC-1-NleE, although effectively halves the RelA activation peak at the start of TNFα addition, does not deplete RelA. After the maximum RelA reduction from 4 h to 5 h post-infection, NleE allows the recovery of RelA signal before the near-complete elimination of cells after 12 h post-infection. The resulting RelA dynamics could lead to reduction in the transient early-stage genes before 5 h post-infection (i.e. within 1 h of TNFα treatment), possibly pro-survival genes to enhance cell death, while having little effect on late-stage gene expression after 5 h post-infection which we have shown to be required for the maximal cytotoxicity of Tir+TNFα (Figure 9D). Overall, in the presence of TNFα, EPEC-1-NleE infection resulted in 95% cell depletion with 70-80% LDH release at 20 h.

High TNFα in PDAC is associated with poor prognosis, while anti-TNFα drugs have been unsuccessful in clinical trials (Egberts et al., 2008; Herman et al., 2013; Padoan et al., 2019; Wu et al., 2013; Zhao et al., 2016). TNFα depletion may instead promote carcinogenesis in certain cases (Waters et al., 2013). The complexity of TNFα in cancer calls for more flexible methods that can tightly regulate the TNFα-NF-κB axis to enhance tumour cell death but not deplete TNFα systematically. From this study, we present these EPEC effector combinations, especially Tir+NleE, as a novel method to efficiently kill PDAC cells, simultaneously limiting the peak of NF-κB activation, in TNFα-rich environment. This method is not limited in simply maximising TNFα-dependent cell death, but capable of fine-tuning NF-κB activation level whilst inducing cell death. The EPEC mutants with individual effectors, effector combinations and effector-TNFα combinations in this study showed a gradient of cell death, similar to those achieved using different Tir dosage and mutations (Zhong et al., 2020). Moreover, they have generated a gradient of RelA nuclear level in PDAC cells (Figure 10). Unlike Tir and Tir_AA_ which do not differ in their RelA regulation activity, the effector-TNFα combinations have different outcomes on NF-κB not correlated to the cell death gradient, presenting an opportunity to simultaneously regulate cell death and NF-κB in opposite ways, and as a result, potentially fine-tune host inflammation and immune responses.

**Figure 10.**
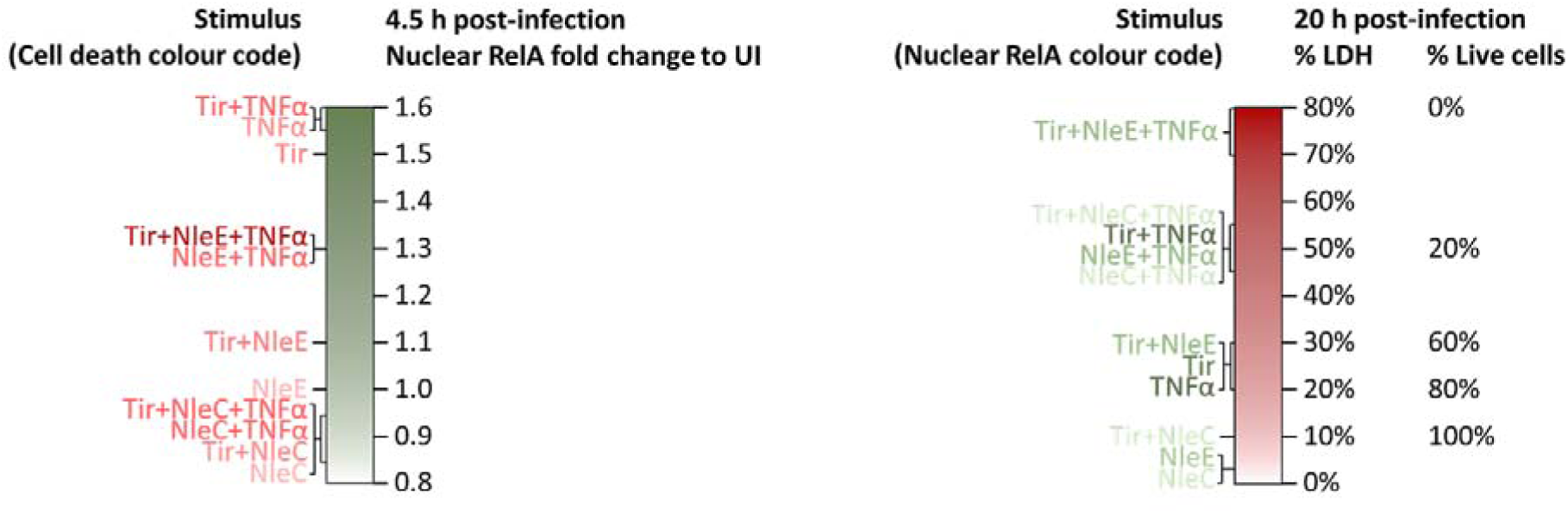
Effector-TNF_α_-induced gradients of RelA and cell death.

Together, these studies showed that the effector functions can be modified using multiple effector combinations and effector-cytokine combinations to manipulate host signalling pathways and engineer highly specific downstream outcomes. We believe that this high level of flexibility of effector-mediated host cell control demonstrated in our work shows great potential for these bacterial proteins to be investigated as future therapeutics.

